# First isolate of *Orientia tsutsugamushi* from Vellore, South India

**DOI:** 10.1101/2023.11.21.568027

**Authors:** Janaki Kumaraswamy, Agilandeeswari Kirubanandan, Lakshmi Surya Nagarajan, Karthik Gunasekaran, KPP Abhilash, John Antony Jude Prakash

## Abstract

**Background:** Scrub typhus a common cause of acute febrile illness in India caused by *Orientia tsutsugamushi* an obligate intracellular bacterium requiring cell culture for isolation. Cell lines like Vero and L929 are most suitable for isolating and maintaining this organism. This study was undertaken to isolate and characterize of *Orientia tsutsugamu*s*hi* from whole blood samples at a tertiary care centre in Southern India.

**Methods:** The PBMCs (peripheral blood mononuclear cells) collected from scrub typhus positive (47kDa qPCR positive) patients were inoculated into Vero and L929 cell line at 80% confluence for primary isolation. The inoculated flasks were incubated at 37°C with 5% CO_2_ for 30 days and examined for presence of *Orientia tsutsugamushi* on the day 10, 15, 20 post-inoculation and everyday thereafter for a maximum of 30 days post inoculation. The scrapings were subjected to Giemsa staining, IFA, 47kDa qPCR and transmission electron microscopy (TEM). The isolates were passaged 3-4 times to ensure viability and then stored in DMEM with 10% FBS (-80°C). Genotyping of the isolates was performed by amplifying a 650 bp segment of the TSA 56 (type specific antigen 56) gene.

**Results:** Amongst the 50 samples inoculated, three were culture positive as confirmed by 47 kDa qPCR at 24^th^ day of inoculation. This was further confirmed by Giemsa, IFA staining and TEM. The 650bp amplicons showed 99.5 to 100% homology with *Orientia tsutsugamushi* MW604716, MH003839, MW604718, MW604717, MH922787 and MH003838 strains. Phylogenetic analysis revealed that 2 isolates belong to TA763 genotype and one belongs to Gilliam genotype.

**Conclusions:** We have successfully isolated and characterised the *Orientia tsutsugamushi* for the first time at our centre from PBMCs. Based on the partial TSA56 gene sequence our isolates belongs to TA763 and Gilliam genotype. More number of samples are being processed for identifying further isolates followed by genomic analysis.

## Introduction

*Orientia tsutsugamushi* is an obligate intracellular bacteria, the etiological agent of vector borne zoonotic disease Scrub typhus (1). *Orientia tsutsugamushi* strains are classified into several subgroups based on the antigenic variations of the major surface protein called type specific antigen 56 (TSA 56) (2,3).

The infection is transmitted to humans (accidental dead-end host) by the bite of an infected larval stage of trombiculid mites commonly called as chiggers (1). In humans, O. tsutsugamushi invades the macrophages and vascular endothelial cells, escapes from the phagosomes, and propagates in the cytosol, inducing an acute febrile state. It often causes a deadly illness, if not treated with appropriate antibiotics, but the molecular pathogenicity is still poorly understood and no vaccine is currently available (4,5).

The scrub typhus diagnosis are mostly based on serological techniques. Currently recognized serological reference standard test is the indirect immunofluorescence assay (IFA) (6). The scrub typhus IgM ELISA also has a similar sensitivity and specificity to the IFA. This assay utilizes the *O. tsutsugamushi*-specific recombinant 56-kDa antigen (7,8). PCR, especially real-time PCR (qPCR) is most useful for diagnosis in the early stage of scrub typhus infection. The common targets of these assays are the genes encoding the 56-kDa antigen and 47-kDa surface antigen, 16S rRNA and groEL genes (9–12).

Isolation of *O. tsutsugamushi* in cell culture, though technically demanding, is useful for understanding the biology including studying the genomics and determining antimicrobial susceptibility (13). Vero and L929 cell lines have been traditionally used to isolate and propagate *O. tsutsugamushi* (15, 16). Growth of the organism is slow (20-30 days), therefore the results are of little use to the patient as well as the treating clinician (14,15). However, it is important to obtain isolates of this bacteria to study the biology of the organisms including pathogenesis, virulence factors, host cell interactions, and genomic features. Further the availability of isolates will be the source of nucleic acids which can be used as positive control for various PCR assays, whole genome sequencing (15-17). The culture isolates can also be useful to develop newer diagnostics and vaccines, which are region specific (15).

Only few studies are available globally on isolation of *Orientia tsutsugamushi* and none from India (Prakash JA personal communication). We, therefore proceeded to establish a standard protocol for isolation, characterisation and maintenance of *Orientia tsutsugamu*s*hi* from whole blood samples at our centre.

## Methodology

### Sample collection and processing

Whole blood (8 ml) was collected in cell processing tubes (CPT) with sodium heparin (BD Vacutainer® CPT™, Franklin Lakes, NJ, USA) from clinically suspected scrub typhus patients. The collected samples were centrifuged at 1600 RCF for 20 minutes at room temperature. The separated plasma was aliquoted and stored; the monocyte layer was pipetted carefully and aliquoted into two tubes with the addition of equal volume of MEM/DMEM. One aliquot of monocytes was immediately transported to cell culture laboratory and inoculated into cell culture; other was used for DNA extraction using the Promega Wizard® Genomic DNA Purification kit (Promega Corporation, Madison, WI, USA) as per the manufacturer’s protocol. The extracted DNA was quantified using Nanodrop Spectrophotometer (ThermoFisher Scientific, Waltham, MA, USA) and stored at -80ºC for molecular assays.

### Isolation of *Orientia tsutsugamushi* in cell culture

The Vero and L929 cell line were maintained according to the protocol described by National Centre for Cell Science (NCCS), Pune. The cells were grown in 25 cm^2^ flasks (Greiner) with growth medium [MEM (minimal essential medium), Himedia, Thane, Maharastra, India] with 10% foetal bovine serum (FBS, Merck, Rahway, NJ, USA) at 37°C and in a humidified atmosphere containing 5% CO2. At 80-100% confluence the cells were sub-cultured into a new flask (1:5) and maintained.

The PBMCs with MEM were directly inoculated into a 10 cm^2^ Vero and L929 cell line at 80% confluence. The inoculated flasks were kept in a shaker for 30 minutes, then at 37°C for 2 hours to facilitate infection. The inoculum was removed and replaced with fresh MEM with 2% FBS and incubated in a 5% CO_2_ incubator. Due to rapid acidification of MEM (Low glucose), the inoculated cell line was subjected to daily media change. To avoid frequent media change we switched to DMEM (with high glucose). For every 3 days once the spent medium was removed and replaced with fresh DMEM contained 2% FBS. The inoculated cell line were observed for morphological changes such as rounding of cells, floating cells, clumping of cells (16). The monolayer was checked by 47 kDa qPCR on the 10^th^ day, 15^th^ day, 20^th^ day and every day thereafter for a maximum of 30 days post-inoculation.After 30 days of negativity the flasks were terminated whereas the positive culture was harvested and sub-cultured into 25cm2 culture flask contained fresh Vero/L929 cell line and maintained in the same way. Giemsa staining, IFA (immunofluorescence assay) and TEM were performed on the *Orientia tsutsugamushi* positive cultures (isolates).

### Bacterial propagation and cryopreservation

The spent medium was removed from infected flasks and replaced with 5ml of fresh DMEM. The infected cell lines were harvested by scraping using a sterile scraper. The detached cells were mixed well and sub-cultured onto fresh Vero cell line at a ratio of 1:5 (1ml inoculum + 4ml DMEM with 2% FBS per flask) and assessed for growth.

**For cryopreservation**, the infected cells were harvested with 5ml of DMEM with 10% FBS and aliquoted into cryovials (1ml). The vials were placed in isopropanol bath [Mr. Frosty freezing container, Thermo Fisher Scientific, Waltham, MA, USA] and kept at -80°C for gradual freezing. After 24 hours, the vials were transferred into liquid nitrogen for long term preservation.

### Giemsa staining

The infected cell line were pelleted by centrifugation at 4000 rpm for 20 minutes. The supernatant was discarded. A loop full of pellet were transferred to sterile glass slide and smear was made. The smear was fixed using methanol and air dried. The air dried smear was flooded with Giemsa Stain, modified solution (Merck, St. Louis, MO, United States) for 20 minutes. The slide was washed with distilled water and observed under light microscope (100X).

### Immuno Fluorescence Assay

Slides were prepared using the above mentioned method. The smear was flooded with primary antibody (IgM ELISA positive serum with OD>2.00) and incubated for 30 minutes at room temperature. The unbound antibody was washed 4 times using wash buffer and air dried. Anti-human FITC labelled antibody was added to the air dried smear and incubated at room temperature for 30 minutes. The unbound conjugate was washed 4 times using wash buffer and air dried. The slide was mounted and observed under fluorescent microscope.

### Transmission electron microscopy (TEM)

The cell culture scrapings were subjected to centrifugation at 4000rpm for 20 minutes. The pellet was pre-fixed with 3% glutaraldehyde and washed in buffer. Then it was post-fixed by 1% osmium tetroxide followed by washing. Then the pellet was blocked with 2% agarose. The block was subjected to dehydration using ascending series graded alcohol (50% to 100%), followed by infiltration by propylene oxide and epoxy resin. Finally, embedded in siliconized rubber mould with epoxy resin and kept in incubator at 60°C for 48 hours for polymerisation. The blocks were subjected to thick section to select the area of interest then subjected to ultrathin section (below 100 nm). The ultra-thin section were taken in copper grid and stained using uranyl acetate and Reynold’s solution which gives contrast. The sections are observed under transmitted to electron microscopy (TEM, *FEI Philips* Technai T12 Spirit, FEI, Hillsboro, OR, USA) and photographed.

### Extraction of DNA from culture isolate

The DNA was extracted from infected cell lines using the Promega Wizard® Genomic DNA Purification kit (Promega Corporation, Madison, WI, USA) as per the manufacturer’s protocol. The extracted DNA was quantified using Nanodrop spectrophotometer (ThermoFisher Scientific, Waltham, MA, USA) and stored at -80ºC for further use.

### Real time PCR

The extracted DNA was subjected to real-time PCR targeting 47kDa gene of *Orientia tsutsugamushi* on an Applied Biosystems™ 7500 Real-Time PCR System (Thermo Fisher Scientific, MA, USA). The amplification was performed using TaqMan Fast Advanced Master mix (Thermo Fisher Scientific, Waltham, MA, USA). Each reaction mixture (25 μl) contained 5 μl of template DNA, 10 pmol of each primer (10 pmol each), 12.5 μL of Master Mix and 5 pmol of the probe. The PCR conditions included an initial denaturation at 95ºC for 5 minutes followed by 40 cycles of 95°C for 30 seconds and 60°C for 1 minute. The Ct value ≤ 35 was considered as positive (17).

### Genotyping of culture isolate

The *Orientia tsutsugamushi* DNA was subjected to a nested PCR protocol as described by Horinouchi. This amplifies a 650 bp segment of the TSA 56 gene (includes VD-1 to VD-III) and was performed as described previously (10,18). The procedure is as given below.

The first round of PCR was performed with 12.5μl of Mastermix, 10pm of outer primers RTS 8 and RTS 9 with 5μl DNA template. The amplicons were generated using the following reaction conditions: 95 C 5 minutes; 35 cycles of 94º C for 50 seconds, 57 º C for 60 seconds, 72 º C for 1.30 minutes and final extension of 72 º C for 7 minutes. The second round of PCR was performed using inner primers RTS 6 and RTS 7 with 2 μl of first round product as DNA template. The cycling conditions are as same as first round except the extension time was reduced to 1 minute. The PCR product was analysed using agarose gel electrophoresis system and visualized by gel documentation (Gel Doc, Bio-Rad, Hercules, California, USA). The 650 bp positive fragment was purified by Wizard® SV Gel and PCR Clean-Up System (Promega Corporation, Madison, WI, USA).

The amplified fragment were subjected to pre-clean up using Exosap-IT Applied Biosystems (ThermoFisher Scientific, Waltham, MA, USA) method with reaction conditions as follows: 37ºC for 15 minutes, 80ºC for 15 minutes and 15 ºC for 2 minutes. The pre-clean up product was used as template for sequencing PCR to amplify individual strands of the fragments BigDye Terminator v3.1 Cycle Sequencing Kit (Applied Biosystems, Foster City, CA, USA). The amplified segment was subjected to post cleanup using HighPrep™ DTR Clean-up System (MagBio Genomics Inc., Gaithersburg, MD, USA) and eluted with molecular grade water. The eluted product was loaded in Genetic Analyzer 3500 (ThermoFisher Scientific, Waltham, MA, USA)] and the raw reads were analysed using Bio-Edit software

### Phylogenetic tree construction

The obtained sequence was subjected to BLAST analysis. Alignment was performed using Clustal Omega (19) with 36 reference sequences retrieved from GenBank-NCBI and the phylogenetic tree was established using IQTREE software: A fast and effective stochastic algorithm for estimating maximum likelihood phylogenies(20)

## Results

Of the 50 samples inoculated into cell culture, Orientia tsutsugamushi was isolated in Vero cells from 3 (confirmed by 47 kDa qPCR) after 24^th^ day of inoculation (refer Table 1). The positive samples (Vellore isolate 1-3) were sub-cultured into 25cm^2^ flask. The three isolates were genotyped by using the 56 kDa type specific antigen partial gene sequence (650 bp). BLAST analysis of this partial sequence showed 99.5 to 100% homology with MW604716, MH003839, MW604718, MW604717, MH922787 and MH003838 *Orientia tsutsugamushi* strains. Phylogenetic analysis revealed that the Vellore isolates 1 and 2 belong to TA763 whereas the Vellore isolate 3 belongs Gilliam genotype (Figure 4).

**Table 1.** Culture isolates positive for 47kDa q PCR.

| S.No | Sample ID | CT value of sample (Monocyte sample) | Ct value of the isolate (Cell line scrapings) |
| --- | --- | --- | --- |
| 1 | Vellore isolate 1 | 26.08 | 18.34 |
| 2 | Vellore isolate 2 | 39.15 | 29.14 |
| 3 | Vellore isolate 3 | UD | 24.43 |

Among the three, 2 isolates were cryopreserved and one isolate was maintained for further morphological analysis such as Giemsa, IFA and TEM (Transmission Electron Microscopy).

Giemsa staining revealed purple colour small rod-shaped organisms in Vero cells Figure 2. IFA showed bright green stained organisms as shown in Figure 3. Further, we analysed the ultrastructural morphology of *Orientia tsutsugamushi* in infected Vero cell by TEM. TEM showed *Orientia tsutsugamushi* in infected Vero cell cytoplasm, the bacterium was round to oval shape with characteristic double membrane (Figure 4B and 4E: indicated by black star), whereas the mitochondria are smaller, darker, spherical/elliptical with cristae seen in higher magnification (Figure 4C and 4D: white arrow head). Extracellular *Orientia tsutsugamushi* are shown in Figure 4F (black arrows).

**Figure 1.**
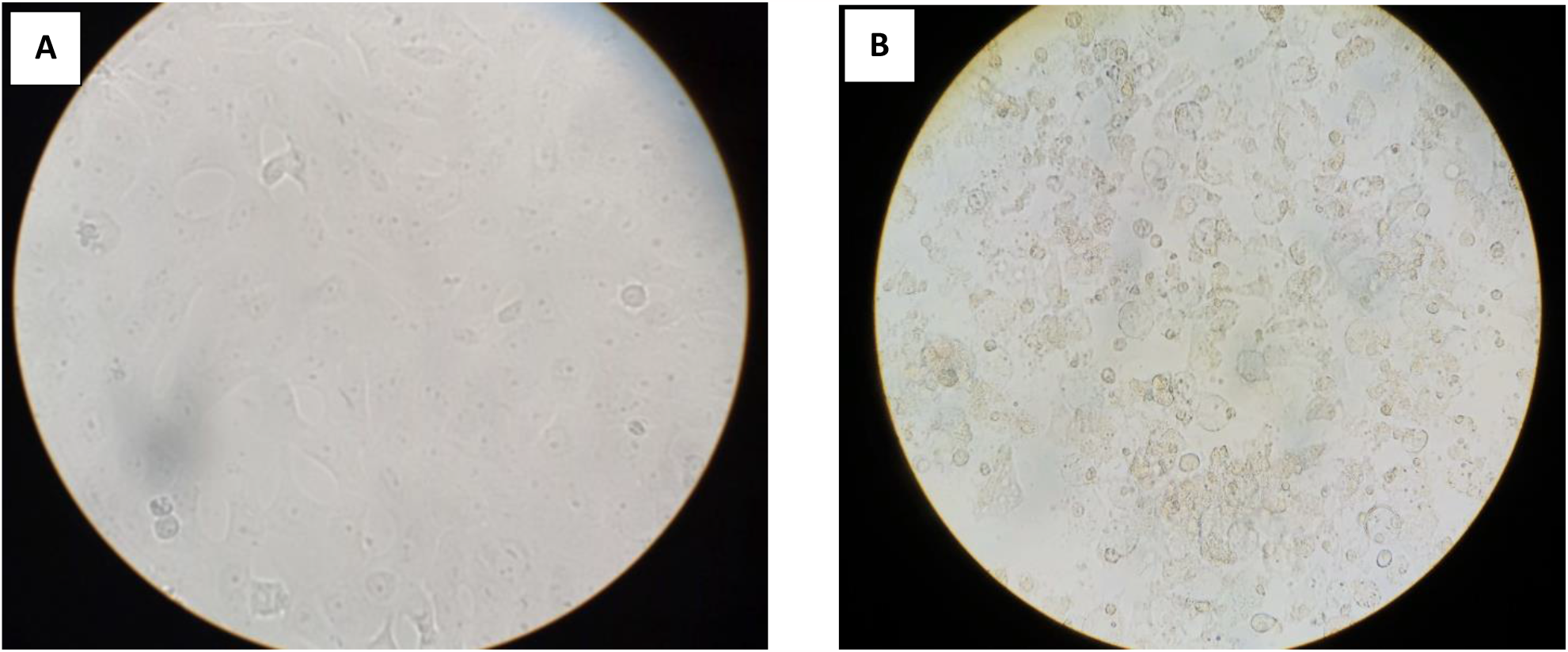
A. Normal Vero cell line, B. Cytopathic effect of Vellore isolate 1 in Vero cell line (Magnification 400X)

**Figure 2:**
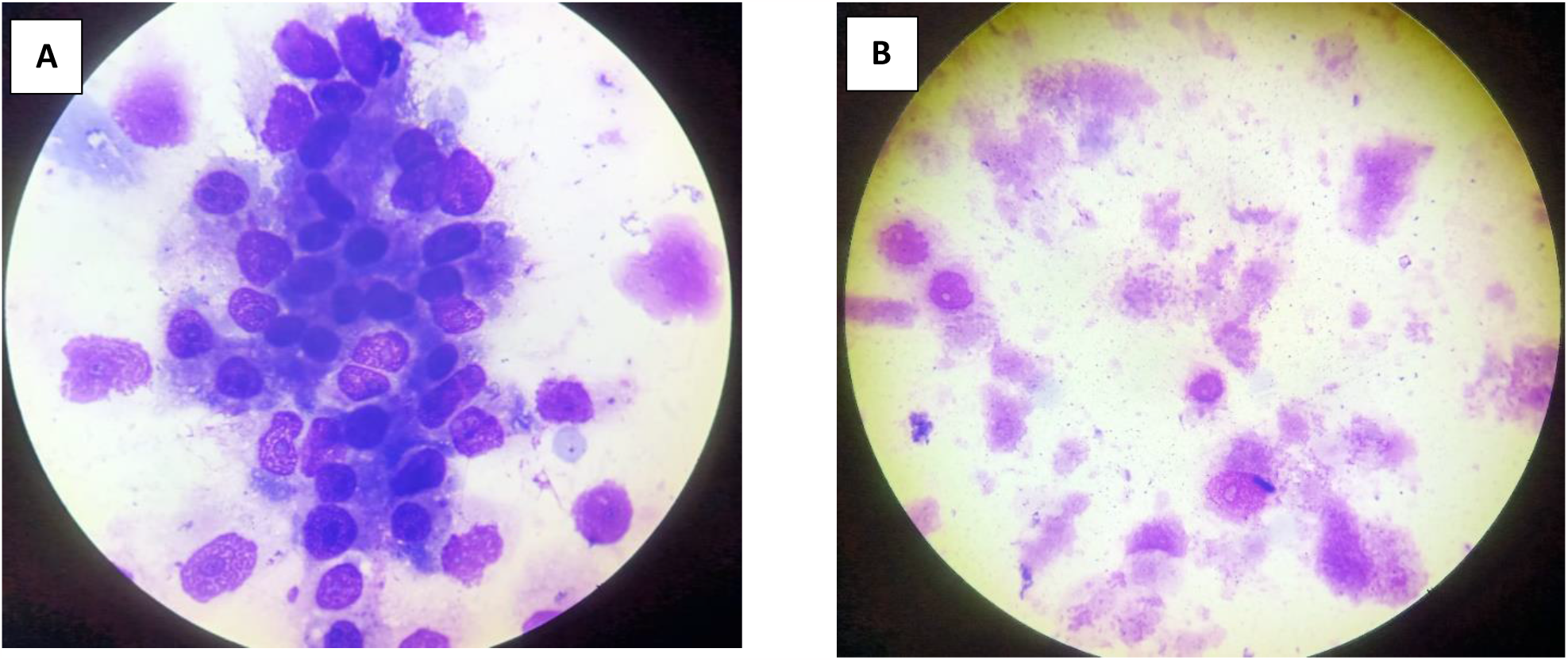
Giemsa staining of Vellore isolate 1 isolate. A. Negative control (uninfected Vero cells) B. *Orientia tsutsugamushi* infected Vero cells

**Figure 3:**
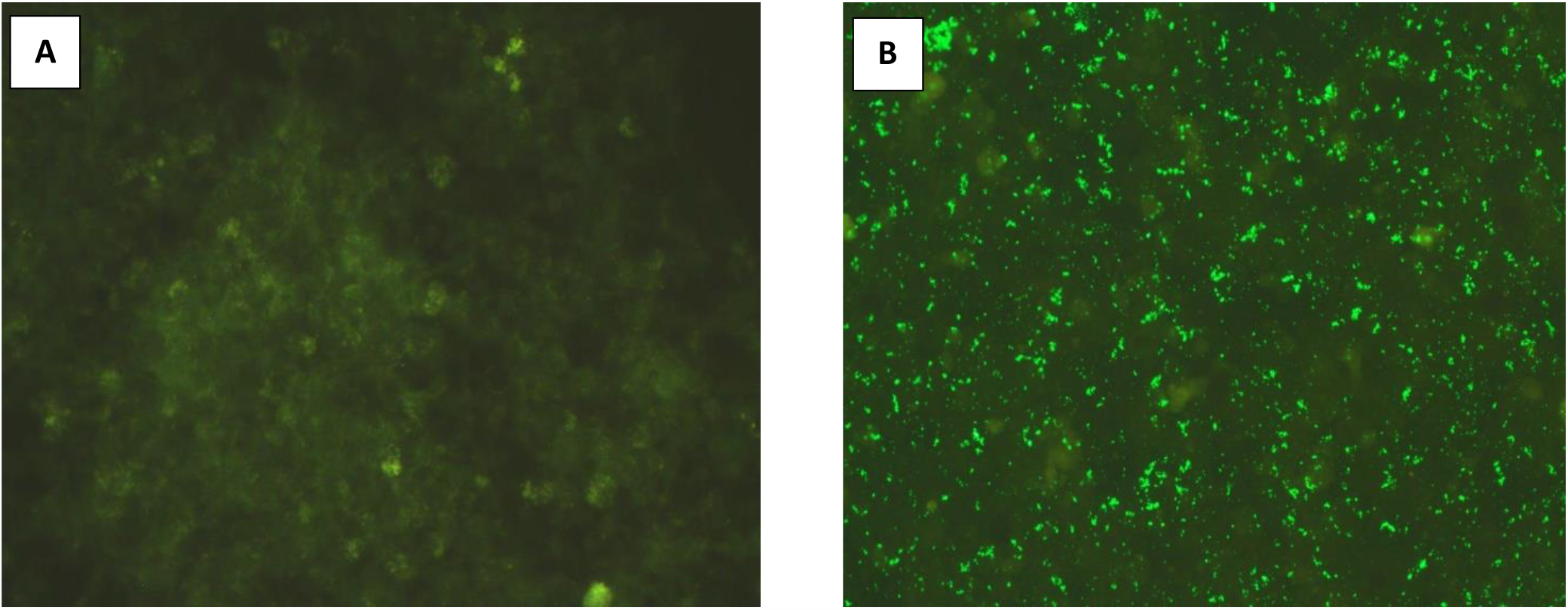
Indirect Fluorescence Assay of Vellore isolate 1 isolate: A. Negative control (unifected Vero cells), B. *Orientia tsutsugamushi* infected Vero cells

**Figure 4.**
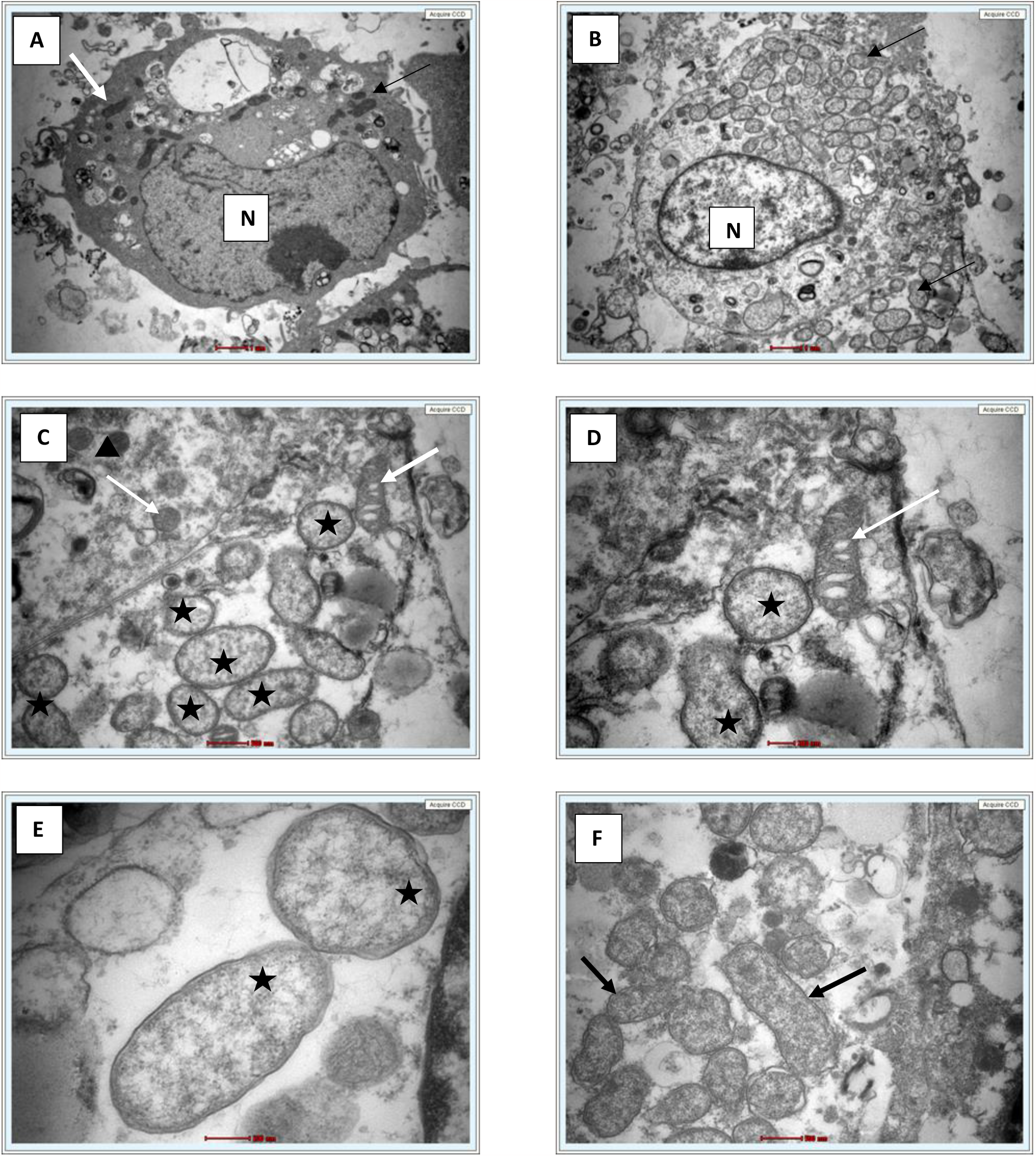
A. Single uninfected Vero cell (6000X) white arrow indicates mitochondria, N-Nucleus. B. Single infected Vero cell (6000X) showed intact cytosolic *Orientia tsutsugamushi* (arrows). C. At 16500X, Intact bacterium (black star) and mitochondria (white arrow), D. At 26500X, bacterium (black star) and mitochondria with cristae (white arrow), E. At 43000X, *Orientia tsutsugamushi* featuring a double membrane and diffuse cytoplasm (black star). F. Extracellular free *Orientia tsutsugamushi* (black arrow) 16500X.

**Figure 5.**
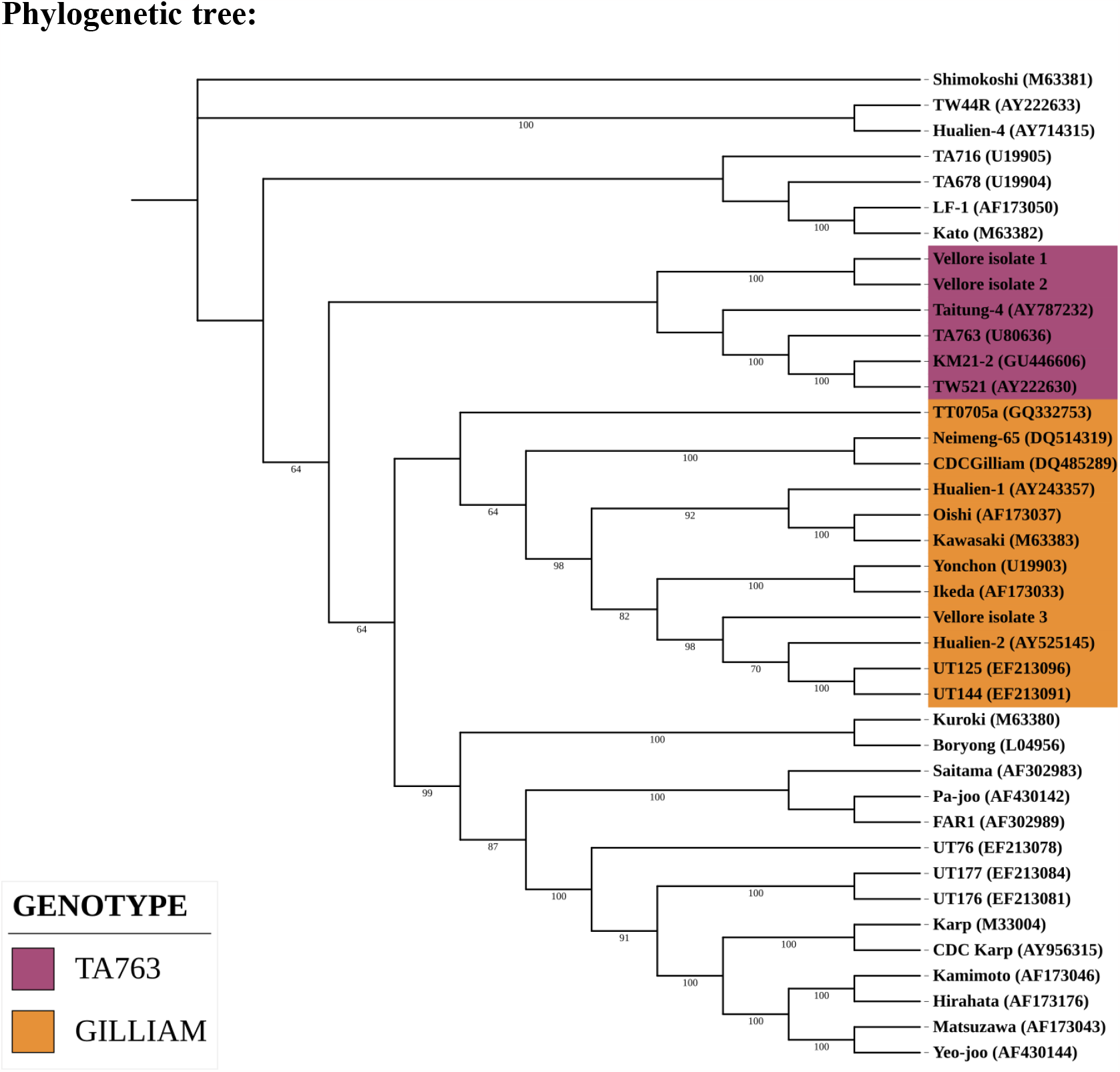
Phylogenetic analysis of Vellore isolates.

## Discussion

The in vitro isolation of *Orientia tsutsugamushi* from clinical samples is important to understand the genomic diversity of the pathogen for development of vaccines and diagnostics. In this study, we have successfully isolated the *Orientia tsutsugamushi* from whole blood sample. The use of 10cm^2^ flask for initial isolation is a simplified method to get an isolate from minimal amount of sample. The 10 cm^2^ culture tube has a wide mouth opening for easy manipulation to sample the infected cell line /supernatant.

Initially, MEM was used for isolation but MEM depleted faster, this required frequent media change. Hence, we switched to DMEM which is more suitable for isolation and maintenance of *Orientia tsutsugamushi* as it has longer incubation period. DMEM is complex media contains four fold increased concentrations of amino acids and vitamins. Cells grow faster and deplete the nutrients slower in DMEM.

Phylogenetic analysis of the partial gene sequence (Horinouchi protocol) which covers the variable domain I, II and III revealed our isolates belongs to TA763 and Gilliam genotype (10,18).

One isolate was morphologically characterised by Giemsa staining, IFA and Electron microscopy. The size and shape of the bacterium resembles mitochondria, the cristae and the difference in electron density helped to identify the *Orientia tsutsugamushi* (Figure 4C and D) as described by Paris and co-workers and cited in the literature by Gemma Vincent (17).

## Conclusion

We have successfully isolated and characterised *Orientia tsutsugamushi* for the first time at our centre and most likely in India from PBMCs. Based on the partial TSA56 gene sequence our isolates belongs to TA763 and Gilliam genotype. Complete 56kDa and whole genome analysis is needed to definitively identify the genotype. More number of samples from suspected cases are being processed for isolation in cell culture to obtain information regarding the genotypes of *Orientia tsutsugamushi* circulating in and around Vellore.

## Author contributions

Conceptualization and methodology: J.K., K.G., A.K.P.P., J.A.J.P.; Investigations and Data acquisition: J.K., A.K., L.S.N.; Analysis and data interpretation: J.K., A.K., L.S.N., J.A.J.P., Drafting the manuscript: J.K., L.S.N.; Critical revision of the manuscript: K.G., A.K.P.P., J.A.J.P.; Supervision and funding acquisition: J.A.J.P.

## Ethics approval

The study was approved by Institution Review Board (Silver, Research and Ethics Committee) of the Christian Medical College (CMC), Vellore, India. **IRB Min. No**. 11942, Dated 27.03.2019

## Acknowledgements

None

